# Cross-sectional white matter microstructure differences in aging and trait mindfulness

**DOI:** 10.1101/301739

**Authors:** Wouter Boekel, Shulan Hsieh

## Abstract

The process of aging can be characterized by a decline in cognitive performance, which may be accompanied by deterioration in specific structural properties of the brain. In this study we sought to investigate to what extent mindfulness changes over the aging process, and which alterations in brain structure can be associated to aging and concomitant changes in mindfulness. We collected Mindful Attention Awareness Scale questionnaire data to assess trait mindfulness and acquired diffusion-weighted imaging data fitted to the diffusion tensor model in a group of 97 middle-aged to elderly participants. Our results showed that trait mindfulness increased with age. In terms of white matter structure our results suggested that there was a general increase of omnidirectional diffusion, which favored radial over axial diffusivity, leading to a decrease in fractional anisotropy (FA) in older participants. We further showed that trait mindfulness mediated the FA-age effect in a localized area consisting of the internal and external capsule, as well as the corona radiata. The implication of this mediation analysis is that trait mindfulness may deter age-associated neurocognitive decline, perhaps by preventing age-associated microlesions specifically in cortico-subcortical white matter tracts. This study can be considered a pioneer of using DTI studies to investigate the relationship between age and trait mindfulness.

**Disclosure statement:** Conflict of Interest: The authors have no actual or potential conflicts of interest.

## Introduction

Healthy aging has become an important research topic, due to increasingly effective healthcare keeping our elders alive longer. Life expectancy has been increasing since the industrial revolution and is approaching an average of 70 years worldwide [1]. Birthrates have gone down in many countries, especially in those with low child mortality rates [1]. In the forthcoming years, wealthy countries will see larger cohorts of elderly people as compared to younger individuals. This shift in the population’s age distribution increases the importance of neuropsychological research aimed at finding efficient ways of maintaining mental health in aging.

Cognitive decline is generally accepted as a “normal” part of aging. Yet, not all cognitive processes will decline with age. So far, research has shown that some cognitive processes (e.g., vocabulary, world knowledge) are less impaired while other cognitive processes such as speed of processing, working memory, and reasoning show large decrements with increasing age [2]. In particular, many studies have provided evidence showing that older adults have deteriorated cognitive control functions, such as less working memory capacity [3,4,5], a deficit in inhibitory processing[6, 7], and a lack of cognitive flexibility [8,9]. One possible fundamental deficit that triggers these cognitive problems with increasing age is a general decrease in the *control* over task-relevant mental processes. That is, it becomes increasingly more difficult for older adults to apply new rules, to coordinate multiple rules, and to maintain relevant information in the context of interfering information.

More practically, age-related declines in cognitive functions have been shown to be associated with severe deficits in the elders’ everyday life activities. For example, some researchers have found performance on cognitive assessments to be associated with instrumental activities of daily living (IADL), such as paying bills, shopping for groceries [10,11]. There are findings showing that updating working memory and task-switching have the strongest relationships with IADL (three domains: home management, financial management, and health and safety). There is also evidence cognitive function influences self-care capacity. For example, learning proper inhaler-use technique has been found to be associated with higher scores on the Executive Interview among older adults [12]. In addition, older adults who exhibited poor performance on tests measuring cognitive flexibility have been found to have increased difficulty in everyday problem solving.

Cognitive function and flexibility have been found to be intimately linked with mindfulness [13,14]. It is important to investigate whether mindfulness plays a role in the aging process. Mindfulness can be divided into two broad categories: state and trait mindfulness [15]. State mindfulness refers to how mindful an individual is in the present moment, hence reflecting a more transient, moment-to-moment experience of mindfulness [16]; whereas trait mindfulness refers to an individual’s natural or innate mindfulness tendency, which has been perceived as a stable, permanent characteristic [17]. Research reporting the relationship between mindfulness and aging has focused more on the manipulation of state mindfulness rather than trait mindfulness. For example, accumulating evidence has suggested that mindfulness-based interventions^1^ (MBI; [18,19,20]) may provide a way to deter the detrimental effects of aging, by promoting body, cognitive, and emotional health [21,22,13,23,24,25].

Mindfulness-based interventions may work because they provide a more holistic attentional training, in the sense that they allow training of general attentional mechanisms, the improvement of which benefits many cognitive tasks. However, neurocognitive research on mindfulness is scarce, and the effect of mindfulness on brain structure has not been elucidated yet. Studies report neural benefits of mindfulness such as increases of overall brain f*ractional anisotropy* (FA) [26] and altered concentrations of important brain metabolites [27] (see Fox et al., 2014 and 2016 [28,29] for reviews on the neurocognitive underpinnings of mindfulness meditation). These findings are interesting, yet there is not enough evidence to allow us to draw general conclusions.

Although research into state mindfulness has important clinical applications, trait mindfulness may provide more insight to infer how individual differences in mindfulness may interact with aging processes [17]. Based on the current literature, it still remains unclear to what extent individual differences in trait mindfulness may interact with the aging process, and whether these interactions are reflected in individual differences in brain structure and plasticity. Prior research discovering differences in the cognitive trajectories of healthy older adults [30] suggests that cognitive decline may not be a necessary consequence of aging, but may result from a range of risk factors that become more prevalent with increasing age. Understanding the mechanisms that contribute to different trajectories of cognitive decline in clinically intact older adults is a significant target for the prevention or reduction in progression of cognitive decline and dementia in old age.

In this study, we investigate whether trait mindfulness shows an association with age and whether brain structural differences (in terms of white matter tract) can inform us about age and mindfulness. In order to answer these questions, we recruited a group of healthy participants from middle-age to elderly and acquired their diffusion-weighted imaging (DWI) data that were further fitted to a diffusion tensor model (i.e., diffusion tensor imaging; DTI) to detect water diffusion directionality, which in turn shows the microstructural architecture of tissue. A questionnaire survey containing the Mindful Attention Awareness Scale (MAAS; [31]) was used to measure trait mindfulness.

## Methods

All procedures performed in studies involving human participants were in accordance with the ethical standards of the institutional research ethical committee at National Cheng Kung University, and with the 1964 Helsinki declaration and its later amendments or comparable ethical standards.

### Participants

A total of 97 participants who were part of a larger lifespan data set collected at National Cheng Kung University were recruited for the present study. All participants were right-handed (Edinburgh Handedness Inventory; [32]), in good general health, without major neurological and psychological disorders, and had normal or corrected-to-normal vision. Demographic information about these 97 participants is presented in Table 1 All participants signed the informed consent form before participating in the experiments. All subjects were paid 1,500 NTD (around $50 USD) after completion of the experiment.

**Table 1.** Demographics information of pariticipants.

|  | Mean | Range | SE |
| --- | --- | --- | --- |
| **Age** | 57.26 | 40-77 | 0.95 |
| **Education** | 13.72 | 4-18 | 0.30 |
| **MoCA** | 26.80 | 22-30 | 0.19 |
| **BDI-II** | 5.12 | 0-13 | 0.42 |
| **PSQI** | 6.16 | 1-14 | 0.35 |
| **MAAS** | 61.48 | 41-90 | 1.15 |

*Note.* SE: standard error; MoCA: Montreal Cognitive Assessment; BDI-II: Beck Depression Inventory II; PSQI: Pittsburgh Sleep Quality Index; MAAS: Mindful Attention Awareness Scale.

### Questionnaires

A set of neuropsychological questionnaires was implemented: (1) The Montreal Cognitive Assessment (MoCA) to screen participants for probable dementia [33]. (2) The Beck Depression Inventory (BDI-II) to screen depression [34]. (3) A Chinese version of the Pittsburgh Sleep Quality Index-PSQI to measure the quality and patterns of sleep [35]. (4) The 15-item Mindful Attention Awareness Scale (MAAS) to measure trait mindfulness [36]. It took between 5 and 10 minutes to complete all questionnaires.

### Imaging Protocols

Magnetic resonance images (MRI) were acquired using a GE MR750 3T scanner (GE Healthcare, Waukesha, WI, USA) in the Mind Research Imaging (MRI) center of National Cheng Kung University. High resolution structural images were acquired using fast-SPGR, consisting 166 axial slices (TR/TE/flip angle, 7.6 ms/3.3 ms/12°; field of view (FOV), 22.4 × 22.4 cm2; matrices, 224 × 224; slice thickness, 1 mm), and the entire process lasted for 218 s. Diffusion weighted imaging (DWI) were obtained with a spin-echo-echo planar sequence (TR/TE = 5500 ms/minimum, 50 directions with b= 1000 s/mm2, 100 × 100 matrices, slice thickness = 2.5 mm, voxel size = 2.5 × 2.5 × 2.5 mm, number of slices = 50, FOV = 25 cm, NEX = 3). 6 non-diffusion-weighted (b = 0 s/mm^2^) volumes were acquired, 3 of which were acquired with reversed phase encoding so as to allow correction for susceptibility induced distortions.

### DTI processing

All DWI data processing and analyses were carried out using FMRIB’s Software Library (FSL, version 5.0.9; www.fmrib.ox.ac.uk/fsl; [37]). Diffusion weighted images were first converted from DICOM to NIFTI format using the MRIcron’s dcm2nii tool (https://www.nitrc.org/projects/mricron/). TOPUP [37,38] and EDDY [39] were used to clean the DWI images of artifacts caused by susceptibility induced distortions, eddy currents, and head motion. A single image without diffusion weighting (b0; b value = 0 s/mm2) was extracted from the concatenated data, and non-brain tissue was removed using FMRIB’s brain extraction tool [40] to create a brain mask that was used in subsequent analyses. DTIFIT (diffusion tensor imaging fitting model; [41]Behrens et al., 2003) was applied to fit a tensor model at each voxel of the data to derive f*ractional anisotropy* (FA), mean diffusivity (MD), axial diffusivity (AD), and radial diffusivity (RD) measures for further analyses.

In order to perform tract-based investigations into DTI measures we performed tract-based spatial statistics (TBSS; [42]) in FSL, which involves voxel-wise statistical analyses of the DWI data. FA images were slightly eroded and end slices were zeroed in order to remove likely outliers from the diffusion tensor fitting. The images were then nonlinearly aligned to each other and the most representative image was then identified. This target image was subsequently affine transformed to 1mm MNI space. FA images were transformed to 1mm MNI space using a combination of the nonlinear and affine registration. A skeletonization procedure was then performed on the group-mean FA image, the result of which was thresholded at FA > 0.2 to identify areas most likely to belong to white matter tracts of nontrivial size.

### Statistical analyses

Linear regressions were performed in R (version 3.0.2; R Foundation for Statistical Computing, http://www.R-project.org) to test for (1) a linear association between participant’s age and their self-reported MAAS mindfulness score, controlling for gender and MoCA, BDI, and PSQI scores as regressors of no interest; and (2) a linear association between DTI measures (i.e., FA, MD, RD, AD) and age, controlling for gender, and MoCA, BDI, PSQI, and MAAS scores (1 regression per measure). In addition, we performed a mediation analysis predicting age based on FA using MAAS scores as a mediator. For the Bayesian version of these tests we used the BayesFactor (http://bayesfactorpcl.r-forge.r-project.org/) and BayesMed (https://CRAN.R-project.org/package=BayesMed;[43]) toolboxes. For the voxel-wise univariate regressions we used FSL’s randomize function on a model mimicking our whole-brain average regressions, testing for a linear association between FA and MAAS scores, controlling for age, gender, and MoCA, BDI, and PSQI scores.

For both the behavioral and DTI results we report classical frequentist p-values, as well as Bayes factors, which provide a more conservative evaluation of the correlations. For ease of reading, we provide both BF_10_ (Bayes factor for the presence of a correlation) and BF_01_ (Bayes factor for the absence of a correlation). These are inversely related (i.e., BF_10_ = 1/BF_01_ and BF_01_ = 1/BF_10_). Bayes factors may be interpreted as proportional evidence for the presence or absence of an effect. For instance BF_10_ of 5 may be interpreted as the data being 5 times more likely to occur under the alternative hypothesis then under the null-hypothesis. In addition, we can interpret the Bayes factor categorically based on a grouping proposed by Jeffreys [44]. Table 2 shows this evidence categorization for the BF_01_, edited by and taken from Wetzels and Wagenmakers ([45]; Table 1, p. 1060). For a detailed explanation of the Bayesian statistics and the Bayes factor, see [46].

**Table 2.**
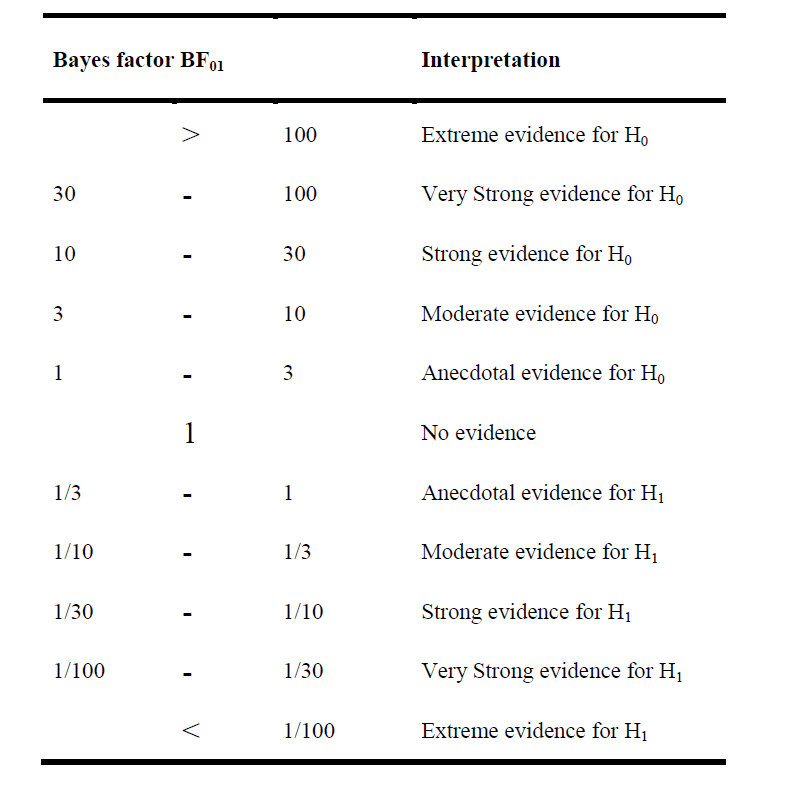
Suggested categories for interpreting Bayes factors based on Jeffreys [44].

## Results

### Age and MAAS

We performed a linear regression predicting age based on MAAS scores. Gender, MoCA, BDI, and PSQI scores were added to the GLM as regressors of no interest. A classical linear regression showed a significant association of MAAS scores (*t* = 3.267, *p* = 0.00154; Fig 1) with age, suggesting that older individuals have higher mindfulness scores than younger individuals (note our sample’s age range: 40 - 77). In addition, a Bayesian linear regression showed strong evidence of a positive association between age and MAAS (BF_10_ = 206.9).

**Fig. 1.**
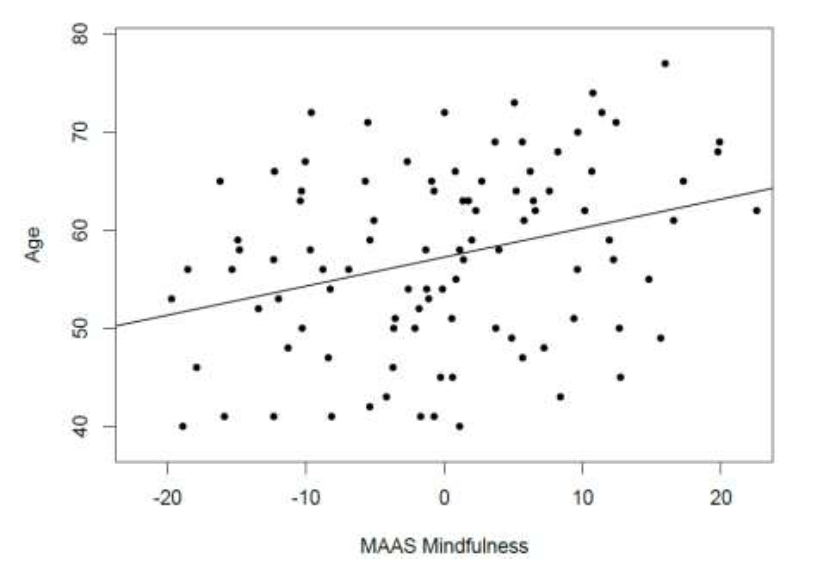
Positive correlation between age and mindfulness assessed by the Mindful Attention Awareness Scale (MAAS). Scatterplot with regression line.

### DTI and Age

We performed a linear regression predicting whole brain average FA based on age while regressing out variance attributed to gender, and scores on MAAS, MoCA, BDI, and PSQI. We observed a negative correlation between FA and age (*t* = −5.497, *p* = 0.000000357, BF_10_ = 21379, Table 3 & Fig 2), suggesting that in older individuals the diffusion tensor is less fractionally anisotropic than in relatively younger individuals. Increases in FA may arise from both a relative decrease in RD, as well as from a relative increase in AD. In addition, MD provides information about the overall omnidirectional diffusion. To elucidate the precise nature of the tensor transformation leading to our observed FA-age relation, we also extracted whole-brain skeleton-averaged RD, AD, and MD, and regressed these against subject’s age (again controlling for gender, MAAS, MoCA, BDI, and PSQI). Results can be seen in Table 3 We observed a positive relation between RD and age (*t* = 5.033, *p* = 0.00000246, BF_10_ = 9540; Fig 3), and non-significant positive relation between AD and age (*t* = 1.895, *p* = 0.0613, BF_10_ = 1.546).For the purpose of understanding the nature of the tensor transformation underlying our observed FA-age effect, it is sufficient to attend to the direction of the correlation, which is positive, indicating an increase in diffusion along the principle diffusion direction (i.e., AD). For the negative FA-age correlation to emerge, the RD has to sufficiently increase as well, which indeed seems to be the case. In addition, we observe a positive correlation between MD and age (t = 4.286, p = 0.0000455, BF_10_ = 754.7; Fig 4), in accordance with the increase in both AD and RD. As such, we may expect that with age there is a general increase of omnidirectional diffusion, which favors radial over axial diffusivity, leading to a decrease in FA. This cross-sectional effect serves as a hypothesis for longitudinal follow-up phases of this study.

**Table 3.** Correlation between FA/MD/RD/AD and Age.

| | *t* | *p* | $BF_{10}$ | $BF_{01}$ |
| --- | --- | --- | --- | --- |
| **FA** | -5.497 | 0.000000357 | 21379 | 0.00004677 |
| **MD** | 4.286 | 0.0000455 | 754.7 | 0.001325 |
| **RD** | 5.033 | 0.00000246 | 9540 | 0.0001047 |
| **AD** | 1.895 | 0.0613 | 1.54641 | 0.6467 |

**Fig. 2.**
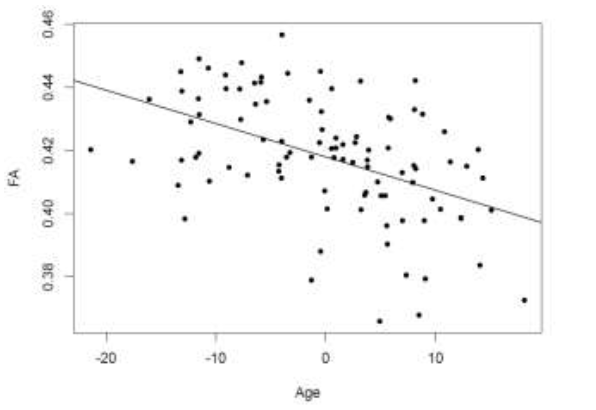
Negative correlation between FA and age. Scatterplot with regression line.

**Fig. 3.**
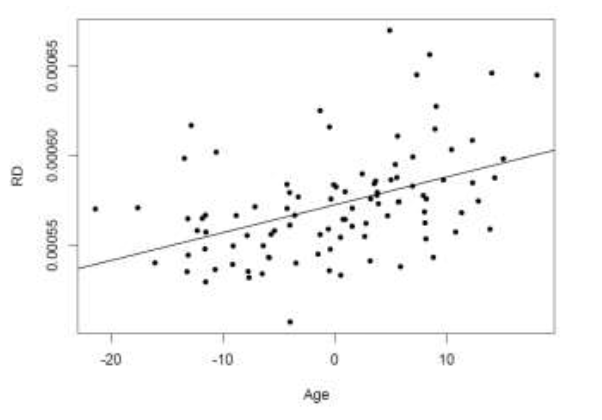
Positive correlation between RD and age. Scatterplot with regression line.

**Fig. 4.**
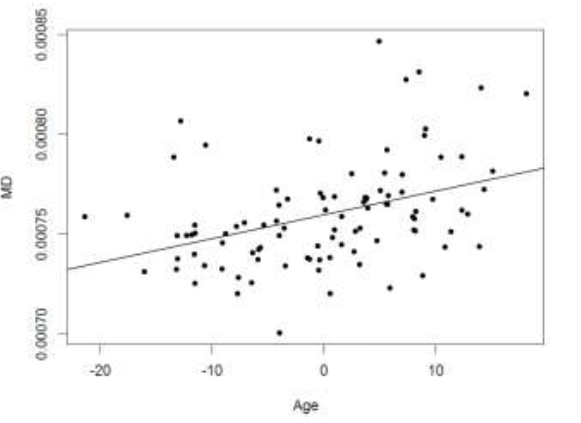
Positive correlation between MD and age. Scatterplot with regression line.

### DTI and MAAS

In addition to cross-sectional age differences, we were interested in cross-sectional differences in DTI measures relating to MAAS scores. We performed a linear regression between DTI measures and MAAS, while regressing out age, gender, and scores on MoCA, BDI, and PSQI. We found no significant relation between MAAS and FA (t = 1.095, p = 0.2764, BF_10_ = 0.2977). Inverting the Bayes factor here provides substantial evidence in favor of the null-hypothesis (the absence of a relation), BF_01_ = 3.576. We subsequently investigated the remaining DTI measures MD, RD, and AD, which were also found not to be associated with MAAS scores (all BF_01_ > 3, Table 4).

**Table 4.** Correlation between FA/MD/RD/AD and MAAS.

| | *t* | *p* | $BF_{10}$ | $BF_{01}$ |
| --- | --- | --- | --- | --- |
| **FA** | 1.095 | 0.2764 | 0.2977 | 3.576 |
| **MD** | 0.074 | 0.9415 | 0.3126 | 3.199 |
| **RD** | -0.311 | 0.7563 | 0.3281 | 3.048 |
| **AD** | 1.026 | 0.3075 | 0.2530 | 3.952 |

We performed an additional mass-univariate regression analysis within the whole-brain white matter skeleton in order to pinpoint spatially localized relations between MAAS and white matter microstructure (i.e., tensor-derived measures). We found a positive relation between FA and MAAS in the left hemisphere, specifically in the internal and external capsule, extending widely from the anterior to the posterior parts of both of these white-matter tracts, and extending dorsally into the corona radiata (*t* = 5.482, p < 0.05 corrected for multiple comparisons; Fig 5). As mentioned previously, differences in FA can be driven by differences in RD as well as AD, so we extracted the mask-average AD and RD from the voxels showing a positive relation between FA and MAAS. We found a positive relation between AD and MAAS scores (t = 3.479, p = 0.000777, BF_10_ = 8.90), and a negative relation between RD and MAAS scores (t = −3.687, p = 0.000387, BF_10_ = 3.02). As such, it seems that in individuals with a higher self-report score on the MAAS mindfulness scale, the diffusion tensor displays a relatively decreased RD, and a relatively increased AD, as compared to low-mindfulness individuals, leading to an overall increase of FA in mindful individuals in the internal and external capsule extending dorsally into the corona radiata.

**Fig. 5.**
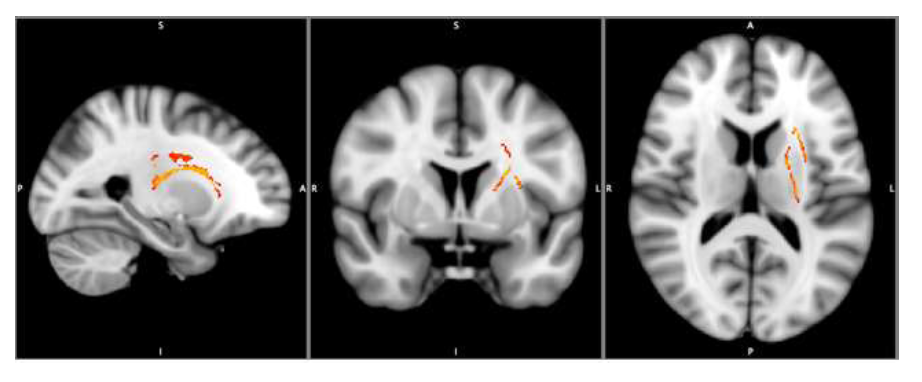
FA-MAAS correlation at x, y, z coordinates: 115, 128, 86.

### Mediation analysis: Age, MAAS, and FA

In light of this finding we were additionally interested in whether MAAS may serve as a mediator between age and FA within the spatially localized area which showed significant MAAS-FA associations. A mediation analysis showed that MAAS was indeed mediating the association between FA and age within this area (BF_10_ = 45.39). We interpret this finding to suggest that as individuals age, FA in this region is preserved, especially for individuals who self-report to be more mindful.

## Discussion

In this study we sought to investigate the relation between characteristics of white matter microstructure, age, and MAAS. To this end we acquired diffusion weighted imaging data in combination with a set of questionnaires (MAAS, MoCA, BDI, PSQI) from a group of elderly participants. We found that age was positively associated with MAAS suggesting that older individuals tend to have higher mindfulness scores. Raes et al. [47] also found a similar age-MAAS effect, as did Mahoney et al. [48], although they used a younger age range and nonlinear aging effects were observed.

In terms of DTI metrics and age our results suggested that in older participants there is a general increase of omnidirectional diffusion, which favors radial over axial diffusivity, leading to a decrease in FA. This is consistent with previous findings which also showed a general decline in FA with age [49,50,51,52].

Our voxelwise investigation into DTI metrics and MAAS found a localized positive association between FA and MAAS in the left hemisphere corona radiata and the internal and external capsule. This finding is in line with some studies using a MBI approach (a measure of state mindfulness), where FA was shown to increase in the corona radiata after intervention. For example, Tang et al [53,54] found that by training participants using a form of mindfulness meditation, integrative body-mind training (IBMT), FA increased in the corona radiata. Corona radiata is an important white matter tract connecting the anterior cingulate cortex (ACC) to other brain structures, so its communication efficiency plays a role in modulating brain activity in the ACC [55]. In line with the finding that FA increased in the corona radiata due to mindfulness mediation, Grant and colleagues [56] observed that cortical thickness in the dorsal ACC was greater for participants with experience in meditation than those without. Therefore, these findings provide convergent evidence for the important role of radiata corona in relation to state mindfulness.

In addition to the corona radiata tract and ACC, some researchers suggest that other brain regions are related to state mindfulness, including insula, temporo-parietal junction, dorsal prefrontal cortex (dPFC), ventro-medial PFC, hippocampus, amygdala, medial PFC, posterior cingulate cortex, insula, and temporo-parietal junction (see [57], Table 2). Based on this neural evidence for an association between brain structures and different facets of mindfulness meditation (e.g., intention, attention, attitude, body awareness, reappraisal, and changes in perspectives on self), Grecucci and colleagues [55] suggest that there are top-down and bottom-up neural circuits involved in the processes of mindful emotion regulation. These circuits include top-down control regions such as the PFC and the ACC and bottom-up emotional regions such as the insula (I) and the amygdala (A). Furthermore, these two neural circuits (PFC-ACC & A-I) may interact with each other via connective neural structures such as the corona radiata. Based on the neuroimaging evidence derived from these MBI studies, the current finding that the FA in the corona radiata tract is positively correlated with MAAS appears to be reconciled with Grecucci et al.’s [55] postulation that the increased FA in the corona radiata tract may facilitate the connection between the two neural circuits (PFC-ACC & A-I), resulting in FA being highly associated with MAAS. Furthermore, our additional mediation analysis showing that MAAS mediated the relationship between age and FA in the radiata corona tract, suggests that as individuals age, FA in the radiata corona is preserved, especially for those individuals with higher trait mindfulness(as reflected by MAAS scores).

It is worth noting that the current finding of a FA-MAAS association is characterized by a thinning and elongation of the diffusion tensor, this is both due to AD increasing with respect to MAAS, as well as RD decreasing. As such, it seems the white-matter in this area is more streamlined for individuals who self-report to be more mindful. The tensor model used here cannot directly inform us about the biological causes of these effects [58], although we could speculate that an overall loss of FA due to white matter lesions occurs with age [51], and that mindfulness in life may prevent some of these lesions. This speculation can be supported by the current mediation analysis showing that MAAS was indeed mediating the association between FA and age within the internal and external capsule, as well as the corona radiata. Future studies could test this hypothesis by employing mindfulness-based interventions longitudinally, since the cross-sectional nature of this study precludes a direct observation of age-related change.

Although the current findings using DTI and trait mindfulness approaches appear to be reconciled with those reported by the studies employing the MBI approach [53,54,55,56,57], research into structure-function associations in trait mindfulness is still considered scarce, and the literature is not yet unified regarding the involved brain systems in state mindfulness. For example, Luders et al. [26] found larger FA in different brain structures to be associated with mindfulness training, including a fiber tract constituting the temporal component of the superior longitudinal fasciculus (tSLF) and a fiber tract linked to the hippocampus, i.e., the uncinated fasciculus (UNC). On the other hand, Tang et al. [53] and Hölzel et al. [57] additionally observed larger FA in an entire fiber tract connecting frontal and tempro-parietal regions, the SLF, and in a fiber bundle linked to the hippocampus, the cingulate cortex hippocampus (CgH), to be associated with active mediation practices. Furthermore, in addition to the aforementioned DTI-based studies, Fox et al. [28] reviewed and used activation likelihood estimation to perform a meta-analysis on the brain structures underpinning mindfulness, in which they observed larger voxel based morphometry in intra- and inter-hemispheric communication (e.g., SLF; corpus callosum) to be associated with meditation. Given these various findings regarding which fiber tracts are related to mindfulness experience, further studies are needed to clarify the involvement of specific brain structures.

## Conclusion

The current results show that with increasing age, an individual’s trait mindfulness tends to increase, but in contrast, whole-brain overall FA decreases due to increasing omnidirectional diffusion. On the other hand, FA in a localized area consisting of the internal and external capsule, as well as the corona radiata, showed an increase with trait mindfulness. Interestingly, we subsequently found that trait mindfulness mediates the FA-age effect in this localized area. Therefore, we speculate that trait mindfulness may deter age-associated neurocognitive decline, perhaps by preventing age-associated microlesions specifically in cortico-subcortical white matter tracts. This study can be considered a pioneer of using DTI studies to investigate the relationship between age and trait mindfulness.

## Acknowledgments

We thank Frini Karayanidis, Birte Forstmann, Alexander Conley, and Wouter Boekel for their great help in setting up this study and Hsing-Hao Lee and Yu-Chi Lin for their help in collecting data. We thank the Mind Research and Imaging Center (MRIC), supported by MOST, at NCKU for consultation and instrument availability.

## Footnotes

1 Please note, in this study, we defined mindfulness-based intervention as a broad term, including a variety of mindfulness-related practices, such as various forms of attention-based and active-based meditations.

